# Bibliometric analysis on cannibalism/infanticide and maternal aggression towards pups in laboratory animals

**DOI:** 10.1101/2023.03.04.531085

**Authors:** José C. Bravo, Lierni Ugartemendia, Arko Barman, Ana B. Rodríguez, José A. Pariente, Rafael Bravo

## Abstract

Animal welfare has evolved during the past decades to improve not only the quality of life of laboratory animals but also the quality and reproducibility of scientific investigations. Bibliometric analysis has become an important tool to complete the current knowledge with academic databases. Our objective was to investigate whether scientific research on cannibalism/infanticide is connected with maternal aggression towards the offspring in laboratory animals. To carry out our research, we performed a specific search for published articles on each concept. Results were analyzed in the opensource environment RStudio with the package Bibliometrix. We obtained 228 and 134 articles for the first search (cannibalism/infanticide) and the second search (maternal aggression towards the pups) respectively. We observed that the interest in infanticide cannibalism started in the 1950s, while researchers started showing interest in maternal aggression towards the pups 30 years later. Our analyses indicated that maternal aggression had better citations in scientific literature. In addition, although our results showed some common features (e.g., oxytocin or medial preoptic area in the brain), we observed a gap between cannibalism/infanticide and maternal aggression towards the pups with only 18 published articles in common for both the searches. Therefore, we recommend researchers to combine both concepts in further investigations in the context of cannibalism for better dissemination and higher impact in laboratory animals’ welfare research.

**Highlights:**

- Cannibalism/Infanticide and maternal aggression have been investigated separately.
- Maternal aggression has a higher impact on scientific literature.
- Combining both topics may increase cannibalism/infanticide impact.

## INTRODUCTION

Early death in mice pups is a common and an accepted problem in laboratory animals’ husbandry and constitutes an undervalued problem due to its extended occurrence. Consequently, there is decreased breeding performance, which has been associated with a reduced welfare in laboratory animals ^1–4^.

Although the mortality rate reported in laboratory mice is high, there is limited knowledge on how pups die. Investigating the different reasons which lead to pups’ death is the key to prevent a lower performance in mice reproduction ^5^. Pups’ death during the first litter is considered normal due to the inexperience of the mother ^6^. However, reproductive efficiency should improve with the experience the females obtain with each successive litter ^7^. Nevertheless, it is not clear yet whether reduced survival rate is related to maternal skills and/or sensitivity to stress. Thus, reproductive efficiency is considered a critical welfare indicator in laboratory rodents ^8–9^.

Maternal care is essential to increase pups’ survival rate and it involves several behaviors toward pups directly or indirectly, including maternal grooming, nursing, nest building or maternal moving among others ^10^. Indeed, during the first 5 days post-partum, spontaneous licks are particularly relevant to maintaining maternal attention ^11^. Maternal aggression is another maternal behavior focused on the defense of the offspring against strange individuals ^12,13^. This aggressive behavior involves both defensive and offensive behaviors, and is provoked by physical and hormonal stimuli ^14,15^. It has been reported that this aggressive behavior decreases over time after the pups’ delivery ^16^. However, on several occasions a mother can show aggressive behaviors towards pups in two different ways. On one hand, cannibalism is the behavior reported in animals that attack and eat intra-specific individuals; on the other hand, infanticide reflects those behaviors which represent killing newborns and pups ^5, 17^. Although both cannibalism and infanticide have the same consequence, they are considered as different behaviors.

Nowadays, bibliometric studies are growing in importance due to their exponential use in research globally. Thus, bibliometric analysis is considered an effective statistical tool for evaluating scientific publications by providing quantitative information about published documents, including patterns, trends or impacts ^18–21^. Moreover, these analyses are able to uncover qualitative information about conceptual relationships, collaborations, thematic maps or relevant keywords to group documents in a given search once they are published. Thus, bibliometric analyses contribute to complete the history of a given field and even highlight the areas of research with lack of studies and, therefore, to propose potential innovations ^22^.

Our topic has been poorly investigated in the previous decades and more information is required to improve successful reproduction in animal facilities. Therefore, our aim was to perform a bibliometric analysis about cannibalism and/or infanticide in laboratory rodents and about maternal aggression towards pups to understand the relationship between both types of research.

## METHODS

### Data collection

Our search was performed on May 1st in 2021 with the following keywords and combined with Boolean (logical) operators:

- To study Infanticide/Cannibalism (I/C):

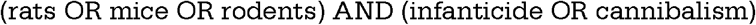
- To study Maternal aggression towards pups (MATP):

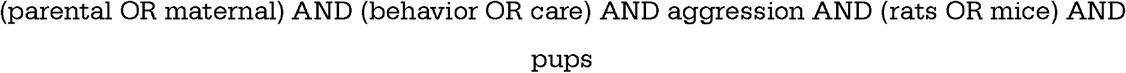

### Bibliometrics analysis of selected manuscripts

The above keyword combinations were used in the Web of Science (WoS, Thompson Reuters, New York). In the first search we found 444 articles. Then, two independent researchers manually selected those articles that fit our objectives and a total of 228 articles were finally included in our study by analyzing the titles and abstracts (see Supplementary File 1 for further details). For the second search, we originally found 267 articles. Following the same criteria explained above, a total of 134 articles were included in the study (Supplementary File 2 for further details). The main cause of exclusion was that some articles were not specifically focused on the topics that we were interested in. The selected data are also available in the GitHub repository https://github.com/rbravo87/Cannibalism_Infanticide_bibliometric_analysis.git Manuscript data were downloaded from WoS and analyzed through RStudio with the Bibliometrix package ^23^. This Bibliometrix package allowed us to analyze and map bibliographic data for a better comprehension of published literature.

## RESULTS

In the context of I/C we observed that 81.5% (186) of the articles were original research studies and 3% (9) were reviews (for more details see supplementary table 1). All the articles found in this search were written in English. We noted that interest in the topic started in 1952 but waned during subsequent years. During the 80s, there was a significant increase (p<0.05; Fig. 1A) in the number of published articles. After the 80s, there was a stabilization for the last 3 decades. However, this trend was not observed in citations: although the average number of article citations per year is stable, the average of total citations per year shows a decreasing trend in the last 20 years (Fig. 1D). The most cited review article we found was by Ebensperger et al., (1998) ^24^ with 203 total citations (Fig. 1B) and the most cited original research article was by Dulac et al., (2014) ^25^ published in *Science* with 172 total citations.

**Figure 1.**
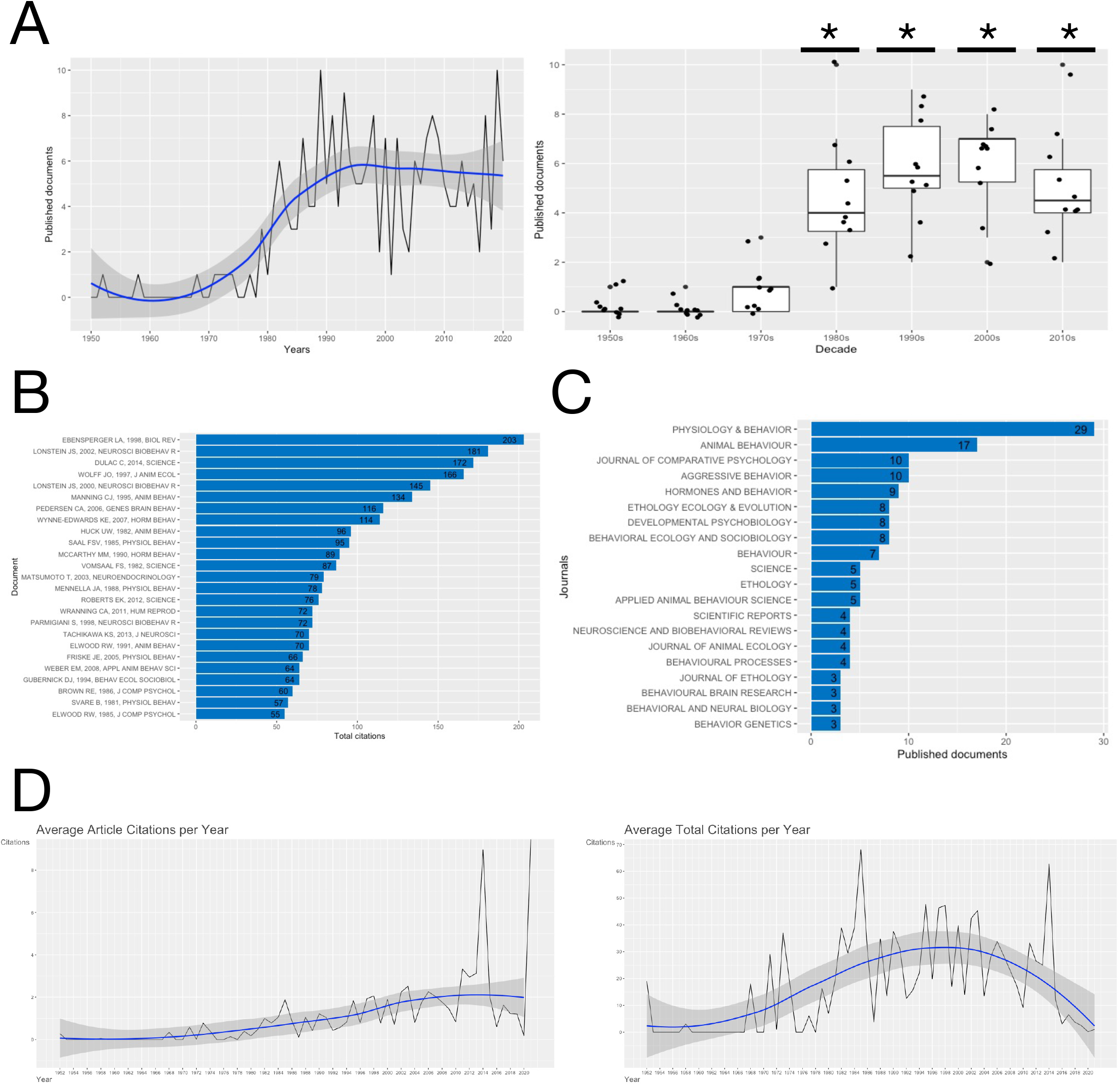
A) Published articles about cannibalism/infanticide in the Web of Science. Left: time evolution. Right: One-way ANOVA comparing publications between different decades, *: p<0.05 when compared with 1950s, 1960s and 1970s. B) Total citations for the top 25 most cited articles. C) Journals which published the articles about cannibalism/infanticide in found in our search. D) Citations per year. Left: Average article citations per year. Right: Average total citations per year.

The articles we found were published in 88 different journals. To have a good idea of the preferred journals, we considered those journals inside the top 25. Our results indicated that *Physiology* & *Behavior* has been the preferred journal since the topic attracted the attention of researchers (Figure 1C). Some years later, *Animal Behavior* led a second group of journals with published documents (Supplementary Figure 1). The top 25 cited articles are shown in Figure 1B. The article with the highest citation was the review published by Ebensperger et al., (1998) ^24^ in *Biological Reviews*. Additionally, researchers from 25 different countries contributed to the publication of retrieved documents. The majority of the articles were published in USA, followed by Italy, Canada and Ireland (Supplementary Figure 2). Finally, when considering the impact of the authors, *Physiology* & *Behavior* appeared again among the top sources and Parmigiani, Palanza and Saal were in the top three of the most relevant authors in our collection for I/C (Supplementary Figure 3).

To study the specific content of the published documents we considered the keywords in the selected articles. In this way, we observed “infanticide” has been the most used word, while cannibalism is not highlighted that much (Fig. 2A). In addition, we observed that the keywords, “maternal behavior” and “maternal aggression” have increased during the past few years in the context of I/C. In general, the most relevant keywords are related to behavior, although we noted some hormones, e.g., prolactin, testosterone, oxytocin, progesterone, estrogen and corticosterone. Further, we observed that cannibalism was the preferred choice of word in 1952 while currently the term infanticide is preferred. Other keywords like maternal behavior have evolved during the past decades to focus more specifically in lactation; however, the term aggression has not changed in the past 5 decades (Fig 2B). Finally, when studying the thematic evolution, we observed that among all the relevant keywords (Fig. 2C), the term infanticide was not used when the interest in the field started. Similarly, during 1997-2010, behavioral studies focusing on maternal behavior became more complex. We observed that the effects of hormones were studied, and currently, the medial preoptic area (MPOA) is considered a key brain region. However, when considering the keywords plus (Fig.2D), the evolution has been simpler and our analysis revealed that progesterone is an essential hormone for the regulation of maternal behavior.

**Figure 2.**
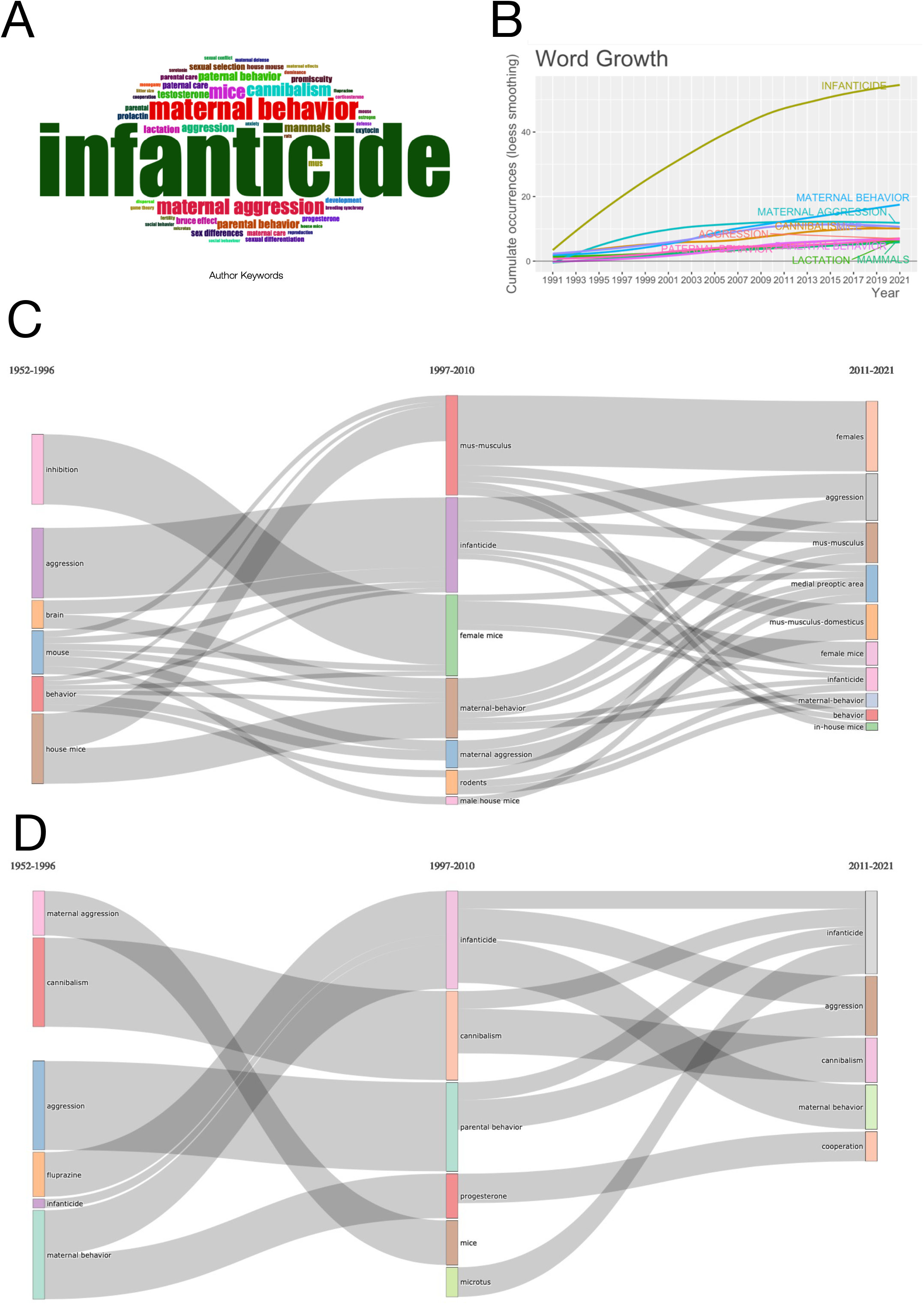
A) Top 50 relevant author keywords in the performed search about cannibalism/infanticide in the Web of Science. B) Keywords word growth in the last 20 years. C) Keywords evolution in three different time slices: 1952-1996, 1997-2010, and 2011-2021.

In our MATP search, we found that 91.7% (123) were original research articles and 5.9% (8) were reviews (Supplementary Table 2) on the topic and all the articles were published in English. The interest in the study of MATP started in 1987 and the scientific research output increased gradually in subsequent years (Figure 3A). When focusing on keywords, we observed that hormones and neurotransmitters have a higher relevance than in the previous topic (I/C). Similarly, mentions of hormones like oxytocin, prolactin, progesterone, corticosterone or vasopressin, and even neurotransmitters like serotonin, nitric oxide or GABA were present (Figure 4A). We observed that the average article citations per year is stable during the past few years, although the average total citations per year showed a decreasing trend (Figure 3D).

**Figure 3.**
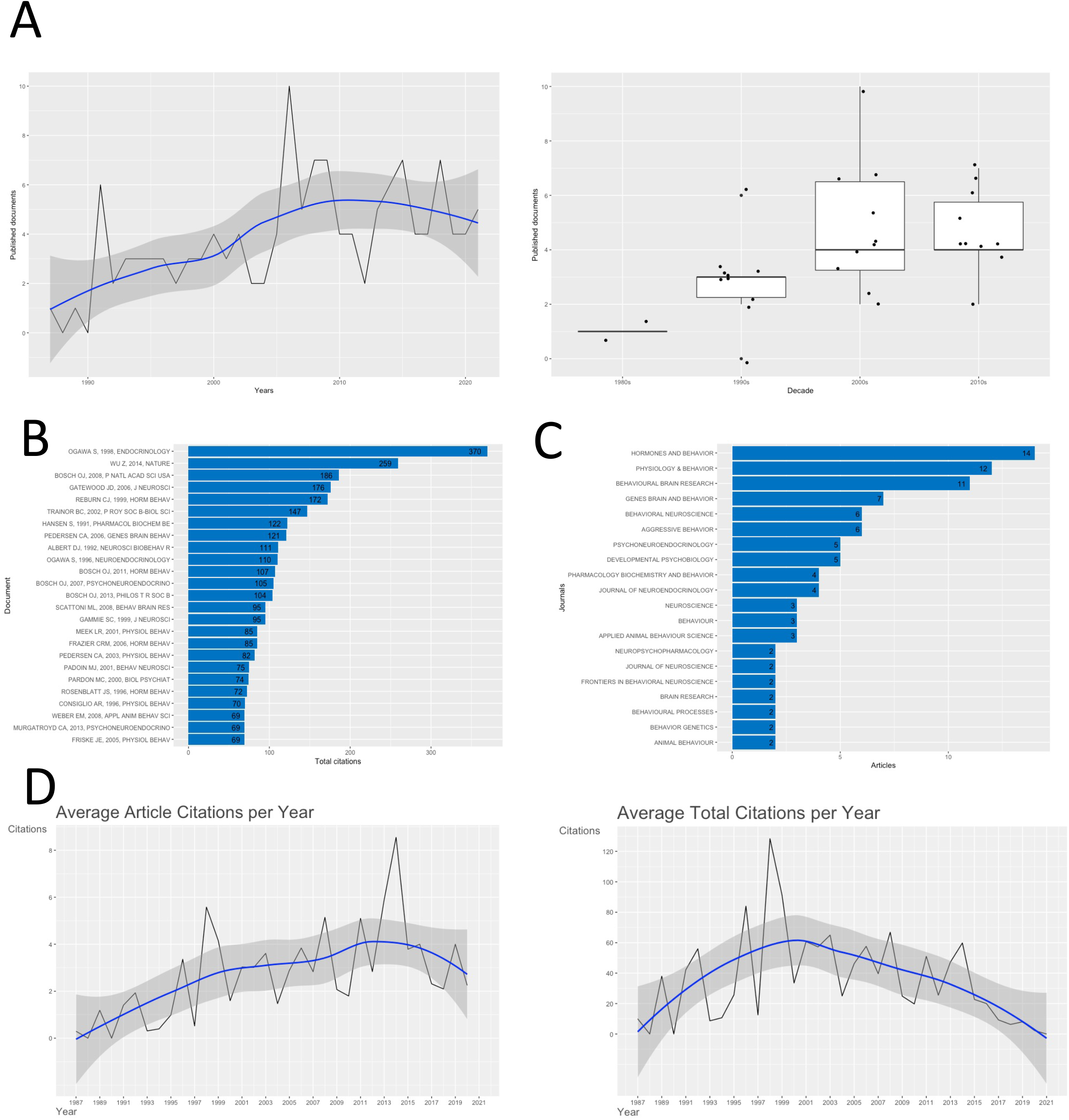
A) Published articles about maternal aggression towards the pups in the Web of Science. Left: time evolution. Right: graphical representation considering the different decades (no statistical analyses are provided). B) Total citations for the top 25 most cited articles. C) Journals which published the articles about maternal aggression towards the pups in found in our search. D) Citations per year. Left: Average article citations per year. Right: Average total citations per year.

**Figure 4.**
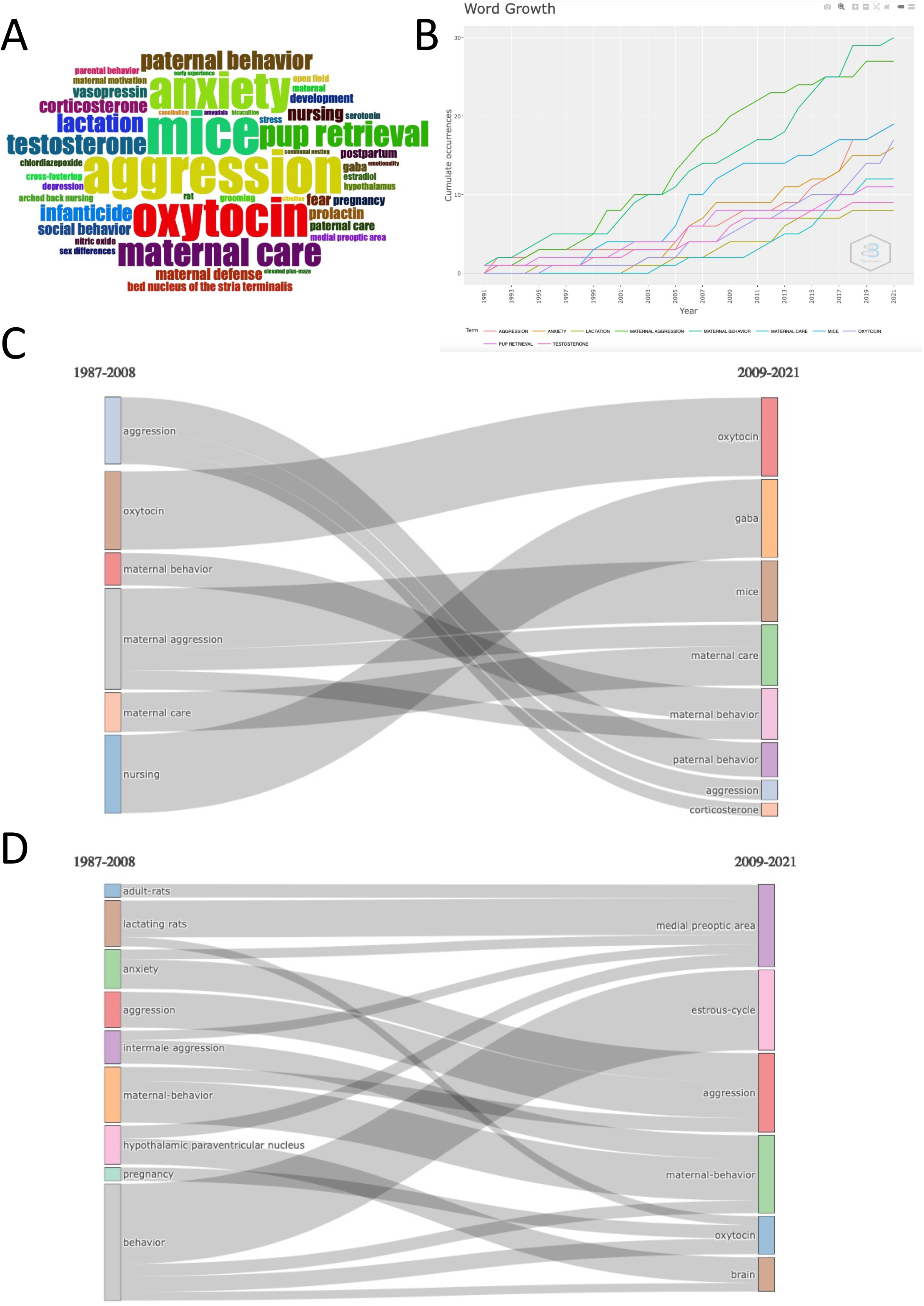
A) Top 50 relevant author keywords in the performed search about maternal aggression towards the pups in the Web of Science. B) Keywords word growth in the last 20 years. C) Keywords evolution in two different time slices: 1987-2008, and 2009-2021.

Articles about MATP were published in 54 different journals. However, three of these journals have published more than half of the publications: *Hormones and Behavior, Physiology* & *Behavior* and *Behavioral Brain Research* (Figure 3C, Supplementary Figure 4). Analyzing the most cited articles, we observed that the top 6 were more cited than the top 6 for I/C (Figures 1B and 2B). The article published by Ogawa et al., (1998) in *Endocrinology* is the one which had the most citations (cited 370 times). Additionally, we found that the top three authors most relevant for the MATP dataset created in our second search were Gammie, Bosch and Stevenson (Supplementary Figure 6).

When we considered the preferred journal to publish research on MATP, we observed that *Physiology* & *Behavior* was the most predominant journal from 2001 to 2015 while *Hormones and Behavior* got more publications from researchers (Supplementary Figure 3). Again, USA appeared as the country with the most number of publications, this time followed by Germany (Supplementary Figure 4).

## DISCUSSION

Bibliometric analyses constitute a relevant tool to understand thematic evolution, and even the connections between the different concepts in a given topic. In this way, this kind of analysis might help in identifying and solving specific problems in different scientific fields ^19^. Moreover, these techniques to analyze scientific literature are able to identify priorities and unify theories ^26,27^.

Animal welfare is a scientific discipline which has seen an increased interest globally in the past few decades, faster than other disciplines and, to date, many variables have been suggested that influence it ^28–32^. Two of the variables widely associated with animal welfare are I/C, which constitute one of the main problems in animal facilities. Therefore, to track the evolution of these concepts for identifying potential areas of interest for further research, we included 228 articles we considered relevant to this topic from 1952 to May 2021. To the best of our knowledge, our research group is the first one in approaching I/C from a bibliometric point of view. Previous research has shown that interest in this field has increased during the past years ^3^. However, we observed that the real interest in this topic started during the 80s, probably due to the principles of the 3Rs (1979). In the same way, Freire and Nicol (2019) ^31^ reported a similar trend during the same decade for animal welfare using a bibliometrics approach. In our search on MATP, we observed that the first publications appeared 3 decades later (during the 80s). Further, we identified that in our search on I/C, *Physiology* & *Behavior* appears as the most preferred journal by researchers, followed by *Animal Behavior*. On the other hand, in our search on maternal aggression, three journals were identified as the main sources with published articles: 1) *Hormones and Behavior*, 2) *Physiology & Behavior* and 3) *Behavioral Brain Research*. Although the topic related to maternal aggression is more recent and has less publications, we noted more citations for these publications (Fig. 2B and 3B).

Interestingly, the evolution of both topics studied in our research show that I/C and MATP have evolved to investigate the role of the MPOA (Figures 2D and 4D). This is likely since the disruption of maternal behavior was established when destroying MPOA in postpartum females by Numan in 1974 and subsequent studies to determine how it is involved in parental behavior ^33,34^. These studies showed that MPOA is a critical structure in the onset and maintenance of parental behavior and further studies will give more importance for maternal care, although a relationship with male behavior also appeared in our analysis. To date, different neurotransmitters and hormones have been proposed to exert important functions for maternal care in MPOA, e.g., prolactin and progesterone, which appear highlighted in our keyword evolution (Fig. 4C) ^35–37^. Thus, prolactin is required for maternal care combined with a decrease in progesterone levels, while aggression seems to be independent of pituitary hormones (and it does not require a decrease in progesterone). In addition, it has been reported that pregnancy, parturition, and maternal care one week after parturition might be controlled by a hormonal stage, and a non-hormonal stage might have a main control starting the second week after the parturition ^38–40^.

In this research, we investigated differences from a bibliometric point of view in I/C and in MATP. We report that both topics have been considered different by researchers and only a small percentage of the published articles considered both perspectives (Supplementary File

3). This is not the first time this kind of a gap is reported, through bibliometric analyses, in animal welfare studies ^41,42^. Further, studies about I/C might increase their visibility due to the observed increase in citations for MATP studies and finally the dissemination of their research will have more impact. Therefore, we encourage researchers who work in the field, to consider both points of view to strengthen this field of study, which will also benefit from a greater interaction between both fields.

## ACKNOLEDWGEMENT

This research was funded by Junta de Extremadura grants (ref. GR21042)

