## Supplementary Figures and Tables for "Bibliometric analysis on cannibalism/infanticide and maternal aggression towards pups in laboratory animals"

Supplementary Table 1: Distribution of the kind of documents published about cannibalism/infanticide

| Documents | Number |
| --- | --- |
| Research articles | 186 |
| Proceedings papers | 7 |
| Notes | 10 |
| Reprints | 1 |
| Early access articles | 2 |
| Editorial material | 2 |
| Meeting abstracts | 11 |
| Reviews | 9 |

ç-

Supplementary Figure 1: Source growth for the articles published about cannibalism/infanticide since 1952.

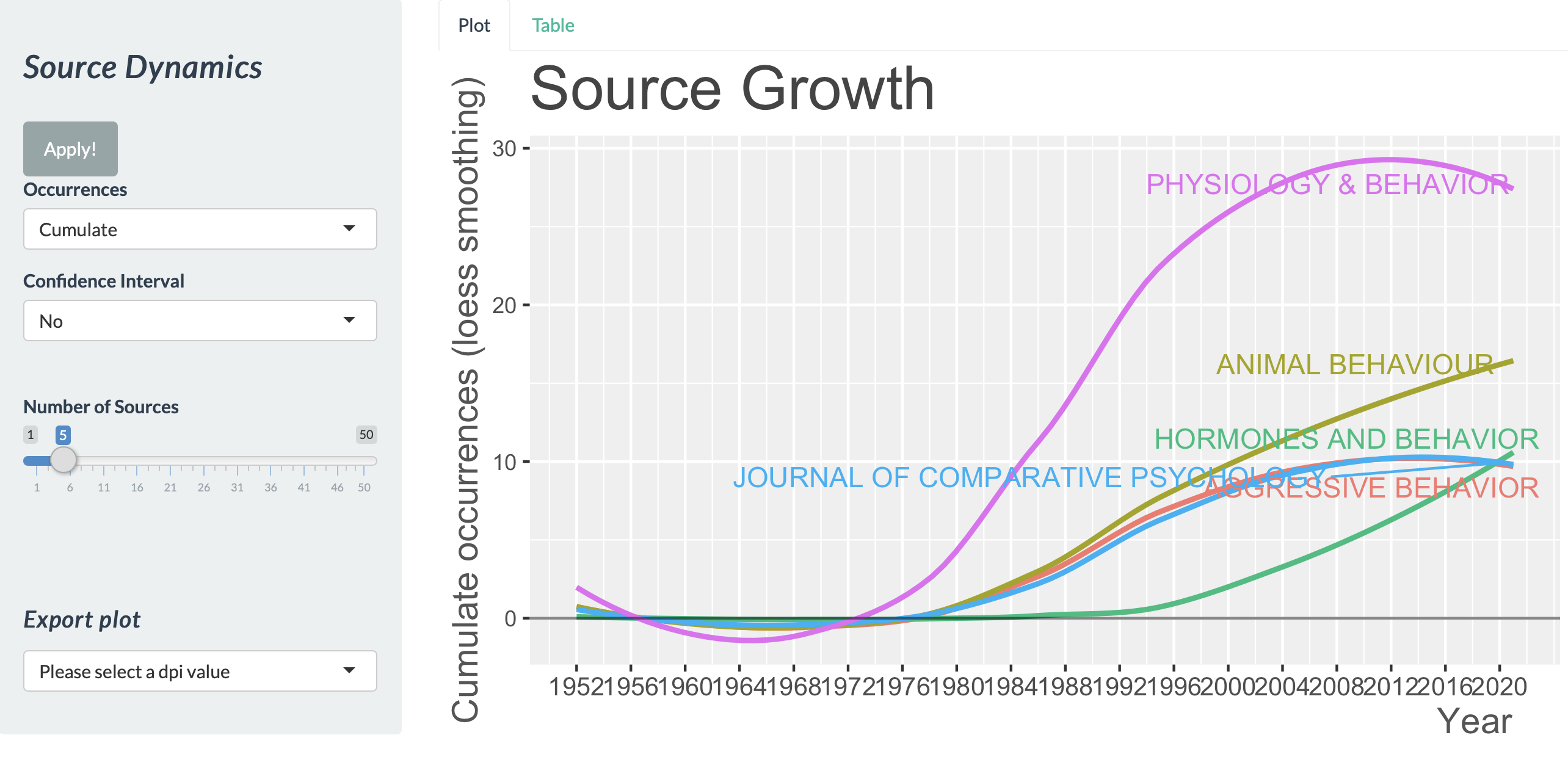

Pie de foto

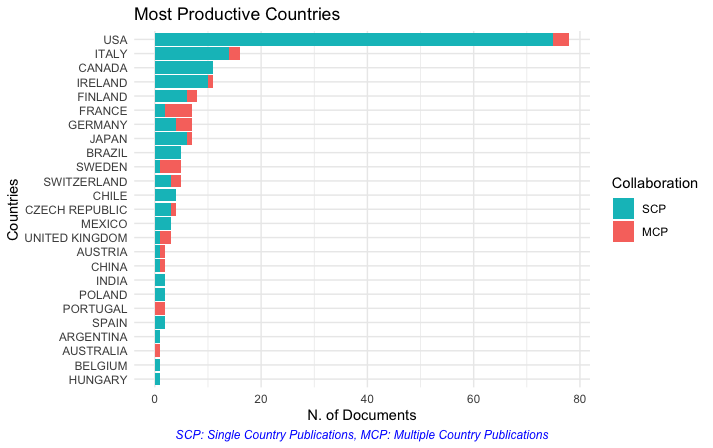

Supplementary Figure 2: Countries contribution to the topic produced by the search about infanticide cannibalism

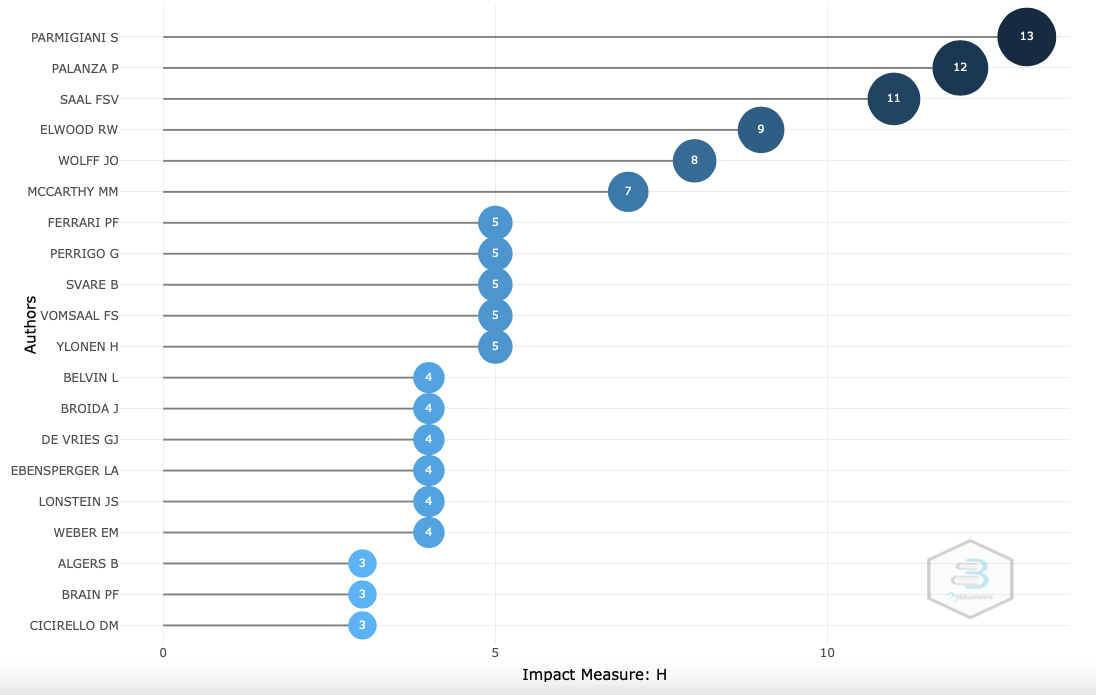

Supplementary Figure 3: 20 authors with more impact in scientific literature in our infanticide/cannibalism dataset.

Supplementary Table 2: Distribution of the kind of documents published about maternal aggression towards the pups

| Documents | Number |
| --- | --- |
| Research articles | 123 |
| Proceedings papers | 2 |
| Meeting abstracts | 1 |
| Reviews | 8 |

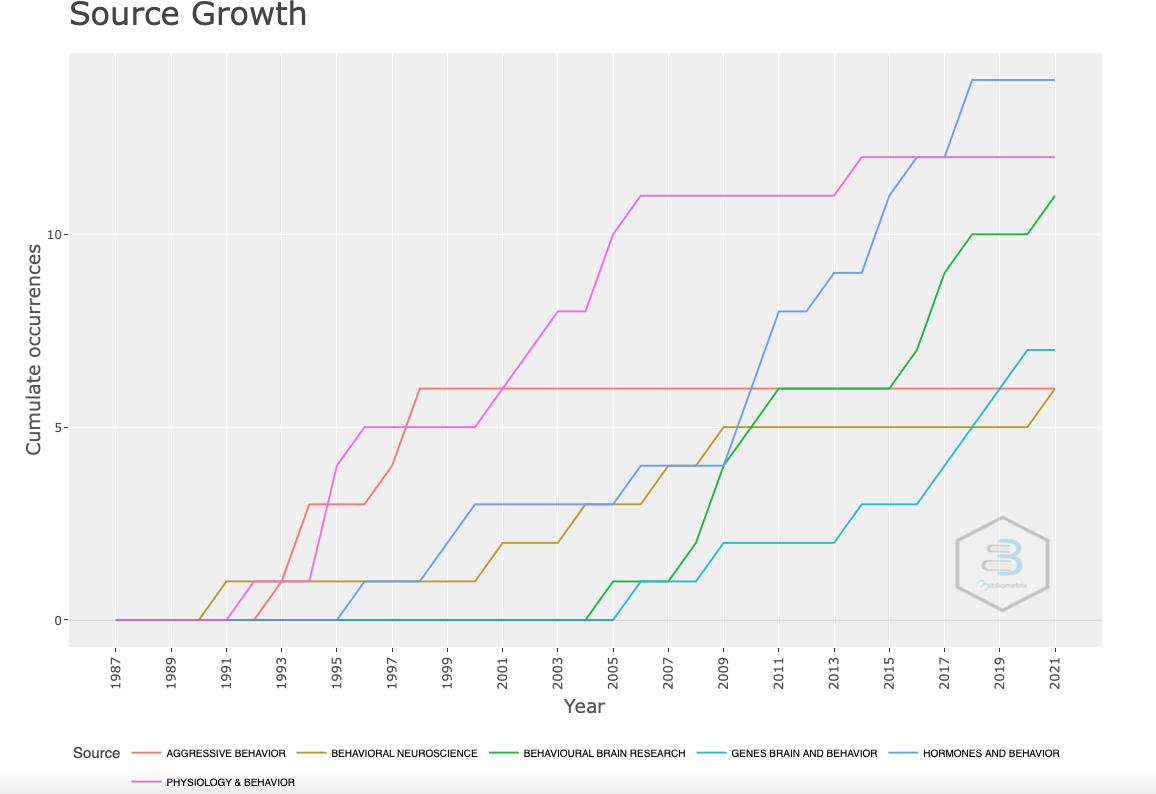

Supplementary Figure 4: Source growth for the articles published about maternal aggression towards the pups since 1987.

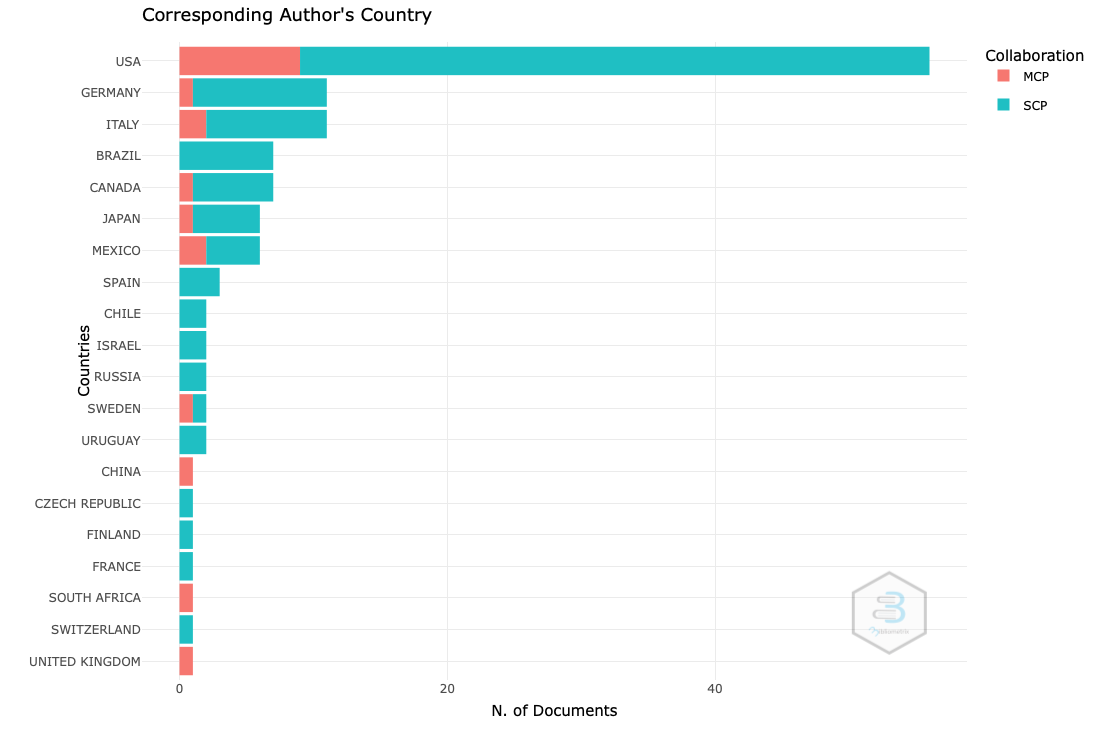

Supplementary Figure 5: number or articles produced by the 20 most productive countries about maternal aggression towards the pups

Supplementary Figure 6
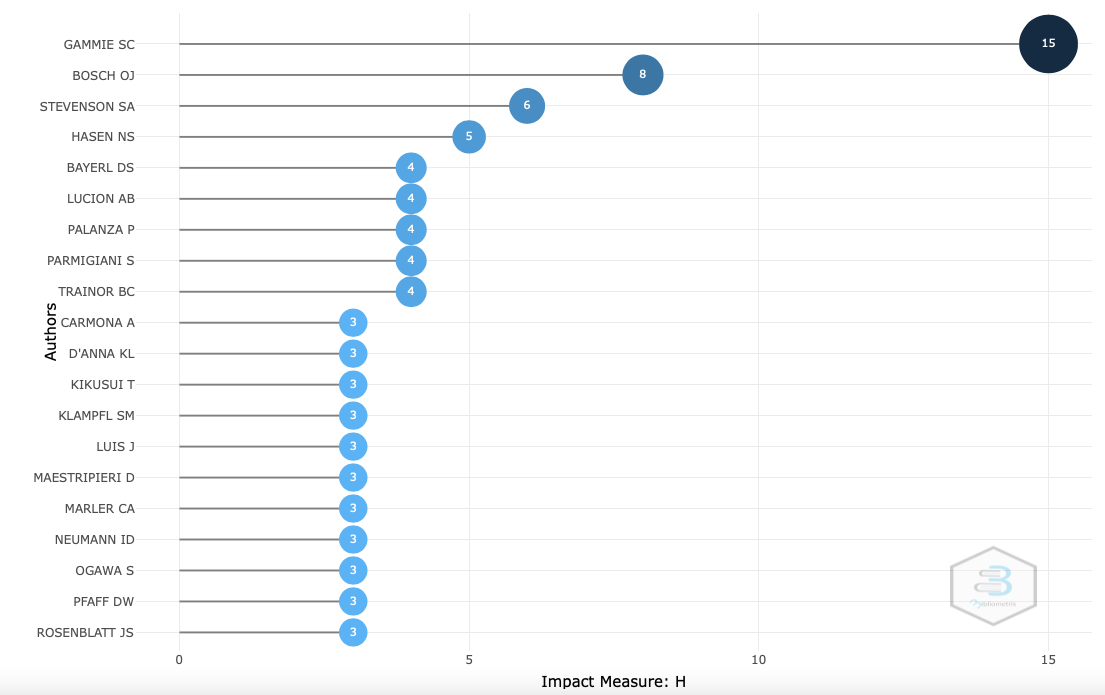
: 20 authors with more impact about maternal aggression towards the pups.
